# Metagenomic and metabolomic analysis of a bryozoan-associated microbial complex from the Hanjiang River, China

**DOI:** 10.64898/2026.09.01.748657

**Authors:** Hui Zhu, Xiaozhi Lin, Ruixuan Wang, Ting Zhang, Wenju Xu, Weifeng Gao, Fang Fang, Jianjian Huang, Yanjie Sun

**Author notes:** **Corresponding:** Ruixuan Wang. Hui Zhu and Xiaozhi Lin contributed equally as first authors to this article.

## Abstract

**Background:** Bryozoans are ancient colonial animals that have existed for over 470 million years and are widely distributed across aquatic environments. Recently, many spherical colloids (average diameter≈1 meter in diameter), containing bryozoans and diverse microorganisms, were found in the Hanjiang river in China. In this study, we investigated these bryozoan-dominated microbial complexes using metagenomic sequencing, metabolomic profiling, and complementary approaches.

**Results:** The results revealed that the microbial structure dominated by bryozoans included eukaryotes such as fungi, microalgae, as well as prokaryotes such as bacteria, and acellular entities such as viruses. All members form a highly integrated consortium that collaboratively maintains community function and stability, which displayed the shape and structure of a regular microbial complex and increased in size. Additionally, the results suggested the predominately presence of Proteobacteria were particularly abundant and are known to produce bryozoans in response to bryozoan signals. Interesting, it was implied that a decrease or even a loss in virulence occurs when pathogenic microorganisms become members of the complex. This is the first study focusing on the composition and metabolites of the unusual microorganism complex. The results discovered the close and intriguing connections within the microbial communities dominated by the ancient organism bryozoan through multi-omics technologies. It showed that both eukaryotes and prokaryotes can coordinate, thereby forming a stable composite structure.

## Introduction

Bryozoans are one of the most ancient metazoan phyla [1]. They are sessile, colonial filter-feeders widely distributed across marine and freshwater habitats and play important ecological roles as benthic community members [2]. They can attach to, or even corrode, many artificial objects (for example, ships and locks) [3,4]. More than 6,000 species have been described globally [5], with many demonstrating rapid growth and reproductive potential, allowing them to colonize new environments and occasionally act as invasive species that alter community structure and biodiversity. However, it has not been elucidated how bryozoans can successfully organize a new microbial complex ecosystem, although they have been originally considered as intruders. Additionally, the disturbance caused by abiotic factors, and the utilization and lack of nutrients in the new environment, are also aspects that need to be investigated in relation to the spreading of bryozoans. Previous data have shown that in nutrient-poor ecosystems, productivity and biodiversity are still maintained [6], mainly because of the presence of many the symbiotic microorganisms. Generally, symbiotic microorganisms include bacteria, eukaryote, archaea, and protozoa [7,8]. It has been revealed that symbiotic microorganisms in sponges are involved with the carbon, nitrogen, and sulfur cycles [7,9]. The ability to fix carbon and nitrogen, and accumulate phosphorus in diazotrophic cyanobacteria that are in symbiosis with sponges could play a central role in shaping the main nutrient flow of the symbiotic microbiota structure [6]. Symbiotic microorganisms, including Rhodobacteraceae, Flavobacteria, Cyanobacteria and other specific taxa present within phytoplankton [10–12] could contribute to the nitrogen fixation [13]. Symbiosis can also lead to competitive exclusion of pathogenic microbes, as scarce resources are exploited [14]. Therefore, the presence of symbiotic microorganisms is very important for the utilization of nutrients in micro-ecosystems, especially within complex aquatic environments. Besides, previous studies have confirmed that microorganisms in symbiosis with bryozoans could produce the bryostatin [15], which could assist bryozoans in predator avoidance and reduction of predator reproduction rate [16]. Moreover, the bryostatin was also shown to be effective in restraining tumors, Alzheimer’s disease, human immunodeficiency virus-1, leukemia, chikungunya virus, ovarian (and some renal) cancers, and the Parkinson’s disease [17,18]. Therefore, it is necessary to study the microbial community of the symbiotic system around bryozoans, their secretions, and the related functions.

In the present study, the recently discovered bryozoan species were identified. Metagenome sequencing and metabolomic analysis were also performed. The aim was to investigate the composition of this unknown microbial complex, and to analyze the role that the entire dominant population associated with bryozoans plays in the microecosystem. The present study will provide an important reference for aquatic ecology research, and will contribute to further studies on the exploitation of aquatic bryozoans to control some important human diseases.

## Materials and methods

### Collection of samples and identification of bryozoans

Spherical microbial complexes (approximately elliptical, with a major axis of ∼31 cm and a minor axis of 16 cm) were collected from the Hanjiang River in Guangdong Province (China), which is an open river and our study did not involve endangered or protected species, and the sampling activity was non-invasive and for scientific research only, so the access was not restricted. From each complex, two types of subsamples were obtained: (i) surface portions with black spots, which consisted mainly of bryozoans, and (ii) the transparent gelatinous matrix. All samples were immediately placed into sterile tubes and transported to the laboratory on ice packs. A portion of each sample was quickly removed with a scalpel for microscopic examination, while the remainder was snap - frozen in liquid nitrogen for metagenomic and metabolomic analyses.Three biological replicates were conducted for all the analyses, synchronously, the black bryozans on the surface of the microbial complex were sampled with tweezer and then were quickly put in the 95% ethanol for species identification, which were labeled. Subsequently, DNA was extracted from the bryozoan sample preserved in 95% ethanol. The target fragment was amplified using a universal primer with the following forward and reverse primer sequences: 5’-GGTCAACAAATCATAAAGATATTGG-3’ and 5’-TAAACTTCAGGGTGACCAAAAAATCA-3’ [19], respectively. Polymerase chain reaction (PCR) was conducted in a 20 μL volume consisting of: 0.5 μL of each forward and reverse primers, 10 μL of 2×PCR Mix premix, 5 μL of DNA template, and 4.5 μL of sterilized distilled water. The amplification program was: 95°C for 5 min, 95°C for 45 s, 50°C for 50 s, 72°C for 50 s, with a final extension phase of 10 min at 72°C, for 35 cycles. The amplification products were examined by agarose electrophoresis, and the target fragments were further purified using a purification kit (Guangzhou Jirui Gene Technology Co., Ltd., China). The amplification products were sequenced also by Guangzhou Jirui Gene Technology Co., Ltd., and the sequencing results were compared with the 18S rDNA sequences obtained by BLAST search in the NCBI database (https://blast.ncbi.nlm.nih.gov/ Blast.cgi). The alignment was performed using ClustalW, and phylogenetic trees were reconstructed using the neighbor-joining algorithm with 1000 bootstrap replicates in MEGA 6.0.

### Isolation and identification of dominant heterotrophic bacteria

Dominant heterotrophic bacteria on the microbial complex were isolated using a sterile ring, and were inoculated on the Brain Heart Infusion (BHI) agar medium plates and thiosulfate citrate bile salts sucrose (TCBS) medium plates. Then, all the plates were incubated at 28°C for 96 h and the dominant colonies were seeded on new medium for purification. After purification, the DNA of the isolates was extracted using the bacterial genome DNA Bacterial miniprep kit (Zymo Research) following the manufacturer’s protocol (Guangzhou Jirui Gene Technology Co., LTD).Then the 16S rDNA gene of the isolates was amplified using the universal primers 5’-AGAGTTTGATCCTGGCTCAG-3’ and 5’-GGTTACCTTGTT ACGACTT-3’ [20](). Then the amplified products were sequenced and the sequencing results were compared with the 16S rDNA sequences obtained by BLAST search in the NCBI database (https://blast.ncbi.nlm.nih.gov/Blast.cgi).

### Sequencing and assembly of microbial genomes on the microbial complex

The total DNA was extracted from samples using the Genome DNA Extraction Kit DNA libraries were constructed using the TruSeq Nano DNA Library Preparation Kit-Set (#FC-121-4001, Illumina, USA) following the manufacturer’s instructions. Metagenome libraries were then sequenced on an Illumina NovaSeq 6000 platform with PE150 at LC-Bio Technology Co., Ltd. (Hangzhou, China). Fastp software (v0.23.4) were used to remove the reads that contained adaptor contamination, low quality bases and undetermined bases.

Then sequence quality was also verified using the Fastp. Quality filtered reads were first aligned to xx genome by using bowtie (v2.2) to filter out host contaminations. Then, the remaining reads were subjected to denovo assembly for each sample using MEGAHIT (v1.2.9) and used to assign microbial functions and taxonomy. Meta Gene Mark (v3.26) was used to predict the coding regions (CDS) of the assembled contigs, and CDS sequences of all samples were clustered using MMseq2 (v15-6f452) to obtain unigenes. DIAMOND (v 0.9.14) was used to perform a taxonomic assessment of the microbiota based on the NR database. The wilcox test was used to identify the differentially abundant species, and significances were declared at P<0.05 and |log2-fold change|>1. An assignment of microbial functions was done using the Kyoto Encyclopedia of Genes and Genomes (KEGG).

### Encyclopedia of Genes and Genomes (KEGG). Gene functional analysis

A clustering analysis (parameters: 95% identity, 90% coverage) was performed using CD-HIT software (http://www.bioinformatics.org/cd-hit/), and the longest gene from each category was selected as a representative sequence to construct a non-redundant gene set. The gene sets were compared with the NR, COG, KEGG, CAZy, CARD, and PHI databases using BLASTP (BLAST Version 2.2.28+, http://blast.ncbi.nlm.nih.gov/Blast.cgi) for biological species annotation and protein function annotation (BLAST comparison parameters were set to an expected e-value of 1e-5).

### Metabolomics analysis

60 mg of bryozoan sample was placed into a 2 mL centrifuge tube and was added with 500 μL of -20°C methanol, 4°C ddH_2_O and 100 mg of glass beads. The contents were vortexed for 30 s. The centrifuge tubes containing the samples were then rapid-frozen in liquid nitrogen for 5 minutes, and thawed at room temperature; they were then placed in a grinder and were shaken at 70 Hz for 2 minutes. This procedure was repeated twice. Subsequently, the samples were centrifuged at 12,000 rpm for 10 min at 4°C. The supernatant was collected, concentrated and dried by rotary evaporation. The samples were then dissolved in 300 μL of 2-chlorophenylalanine (4 ppm) in methanol (1:1, 4°C) and filtered through a 0.22 µm membrane [21]. The samples were subjected to LC-MS detection. The detection conditions were the same as those previously reported by Zhao [22].

### Correlation analysis of metabolites and microbes

To further investigate the relationship between microbial communities and metabolites, a Pearson correlation analysis was performed between the metabolites obtained from metabolic annotation, and the microbial phylum levels obtained during macrogenome sequencing. The criterion for screening was a correlation coefficient |r| > 0.50. Network diagrams were drawn using the R language igraph package (version 1.1.1).

### Data Aavailability

The original contributions presented in the study are included in the manuscript and supplementary material. Data are available from the Ethics Committee of Hanshan Normal University (contact via Hanshan Normal University) for researchers who meet the criteria for access to confidential data.

## Results

### Description of the microbial complex and identification of the dominant bryozoans

The microbial complex had a round or oval shape (Fig. 1A), and resembled an air bag; the texture was like that of jelly. Most microbial complexes reached a diameter of about 60∼100 cm, and could float on the river surface. During sampling, the water temperature was 33.5°C, salinity was 0.06, pH was 8.33∼8.35 (because during the sampling phase, large amounts of algae appeared in the river, which changes the pH of the water environment), dissolved oxygen concentration was 7.89 mg/L, conductivity was 148.6 us/cm, and the water flow rate in the river was 2.83 cm/s. When climatic conditions changed (for example when it rained), this microbial complex could quickly settle to the bottom of the river and live through fixation. The surface of the complex, which presented a fixed and regular texture, was covered by bryozoans, whose population showed an absolute superiority in numbers (Fig. 1B). Observed under a microscope (Fig. 1C), individual bryozoan was disc-shaped, presented a hard texture, and the diameter was 1 to 1.5 mm. The central region of each colony was solid, and around it, there were 16 to 24 hooked spines, showing an asymmetric distribution. All the hooked spines radiated from the annulus periphery, and each one was composed of fused extensions of the dorsal periblast (Fig. 1D). There was a “scaffold” inside the hooked spines, at the top of which there were two pairs of barbate spines growing in opposite directions from each other (Fig. 1E, F).

**Fig. 1.**
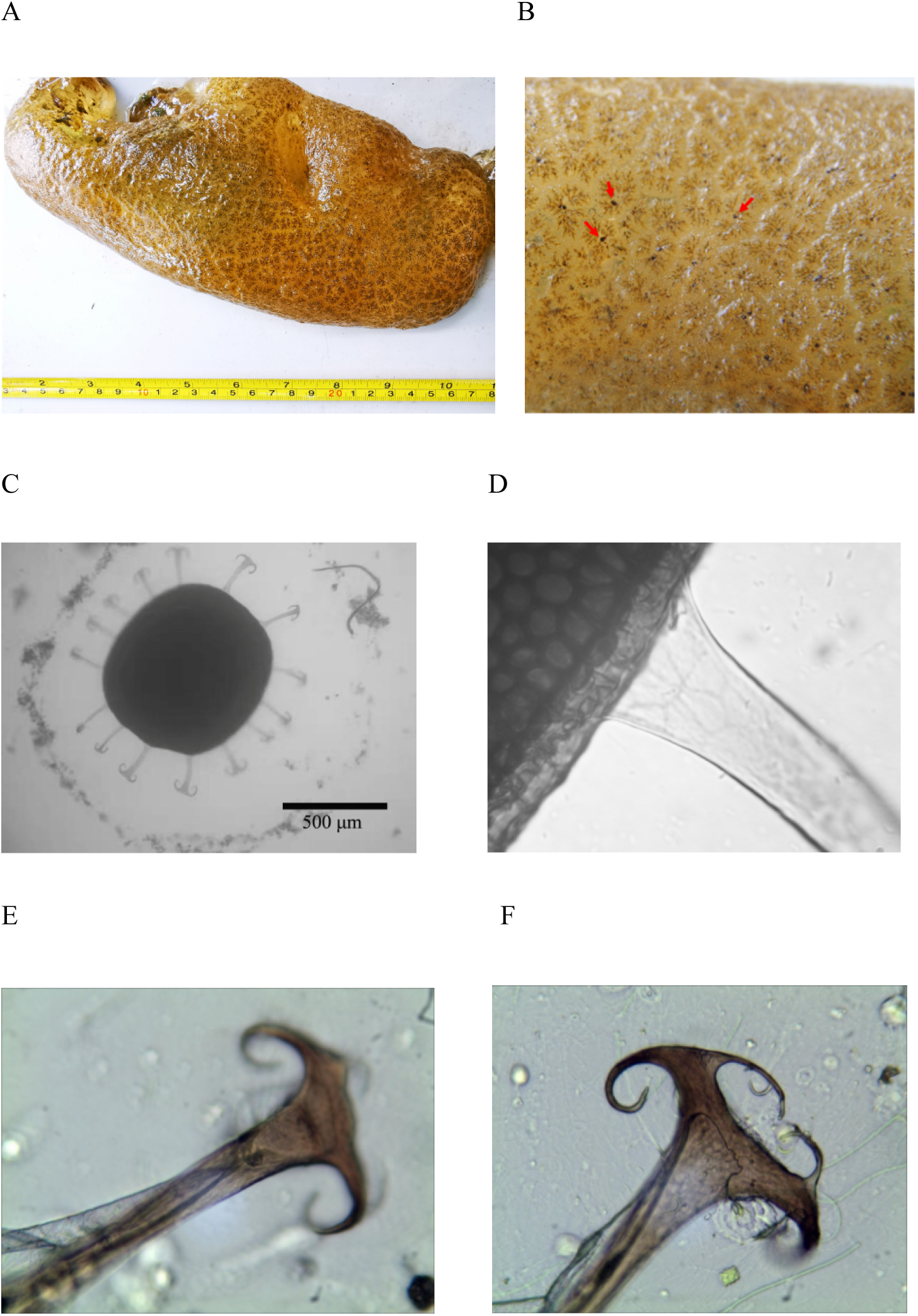
Identification of the bryozoan. **A** Microbial complex. **B** Regular pattern on the surface of the bryozoans. **C** Morphology of the bryozoan under the microscope. **D** Spines of the bryozoan originated from the edge of the ring. **E** Hooked spines. **F** Hooked spines.

The bryozoan sequences have been confirmed and compared with the 18S rDNA sequence in GenBank. The results showed that the bryozoan (labeled as HJBry2020) belonged to *Pectinatella magnifica* (Fig.2). The bacteriological testing of samples revealed that the culturable heterotrophic bacteria on the surface of the microbial complex consisted of the *Pseudomonas* (25%), *Bacillus* (25%), *Photobacterium* (25%), *Acinetobacter* (12.5%), and *Shewanella* (12.5%) (Table S1).

**Fig. 2.**
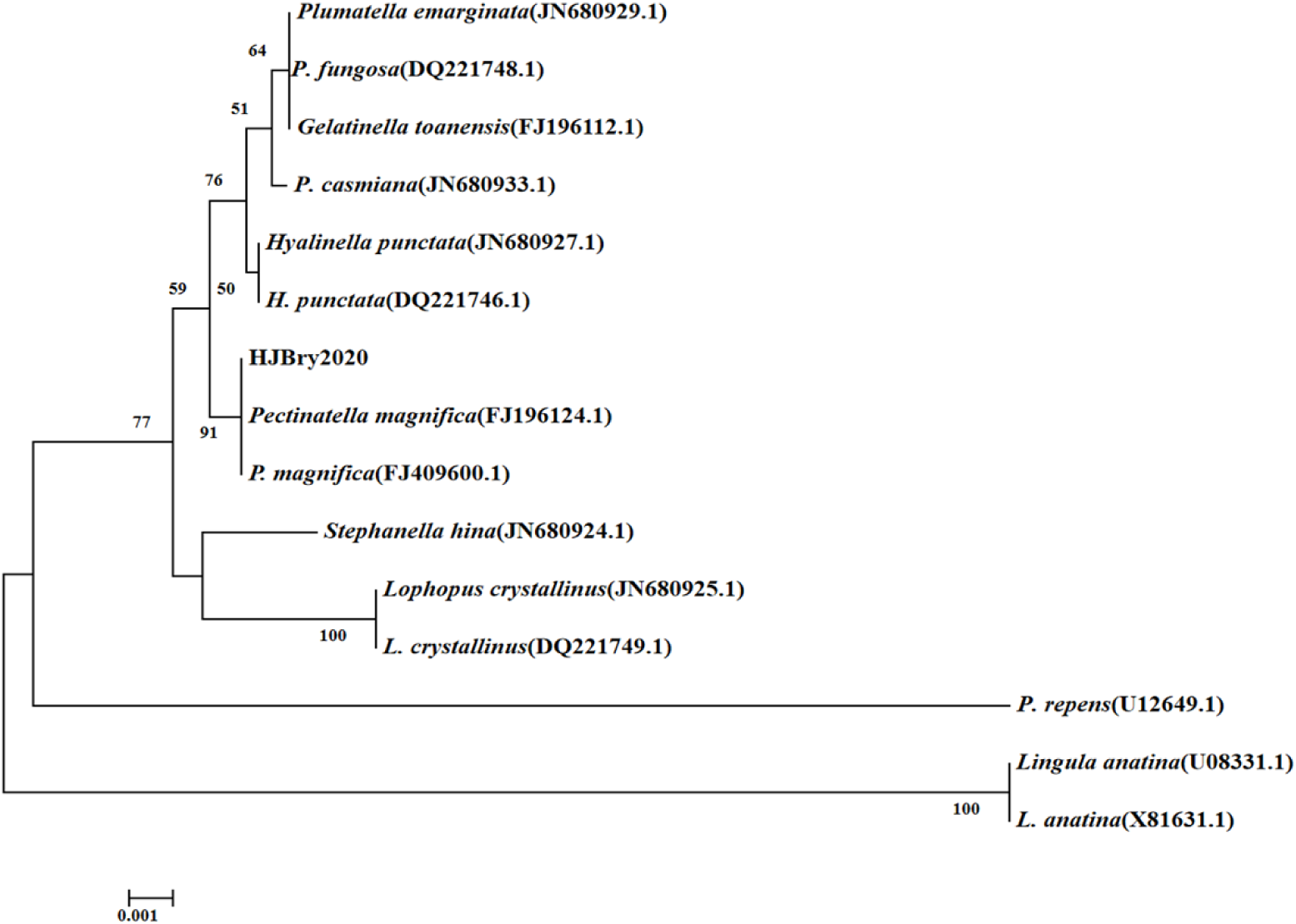
Phylogenetic tree based on the 18S rDNA gene sequence of the HJBry2020.

### Characterization of the microbial genome of the bryozoan-dominated microbial complex

Approximately 54.62 Gbps of high-quality sequences in total, with an average of 18.20G per sample, were generated using an Illumina Hiseq instrument (Table S2). The average assembly size of the metagenome ranged from 597,490,495 bp; the average N50s were 5,004 bp. The average number of open reading frames (ORF) predicted was 87,383. The average GC content was 47.99%.

### The presence of microbial communities contributes to the use of substances and to defend the microbial complex against pathogenic microorganisms

As shown in the Figure 3, bacteria (10.79%) were the most abundant organisms, followed by the fungi (1.04%), viruses (0.05%), and archaea (0.03%). A total of 64 bacterial phyla, including 740 genera, as well as 1285 species were observed (Online Resource 1). Large amounts of Proteobacteria, as well as Bacteroidetes and Actinobacteria, were annotated in the microbial complex, *Acinetobacter* and *Escherichia* were overwhelmingly dominant (Fig. 3). At the species level-and consistent with the genus level-the most abundant organisms were *A. baumannii*, Gammaproteobacteria and *E. coli*, which suggested the absolute dominance of Proteobacteria in the overall microbial community, and a significant number of Bacteroidetes were also found in the microbial complex. Additionally, ten fungal phyla, comprising 389 genera and 650 species, were also detected within the microbial complex samples (Online Resource 2), and Mucoromycota, Chytridiomycota, and Ascomycota were the most dominant, including the *Gonapodya* and *Synchytrium*. The top three species were *Rhizophagus irregularis*, *G. prolifera*, and *S. microbalum* (Fig. 3). Moreover, a small number of viruses was also present in the complex (accounting for 0.05%), and 47 genera and 83 species were observed (Online Resource 3). The predominant viral genera were *Alpharetrovirus*, *Errantivirus* and *Gammaretrovirus*, while the top three viral species were *Avian sarcoma virus*, *Lampyris noctiluca errantivirus* and *A. leukosis virus*. The presence of archaea in the microbial complex was also detected, including Euryarchaeota, Bathyarchaeota and Thaumarchaeota, which were the most abundant archaeal phyla (Fig. 3). The three most abundant archaeal groups at the order level were Methanosarcinales, and at the phylum level were Bathyarchaeota and Euryarchaeota.

**Fig. 3.**
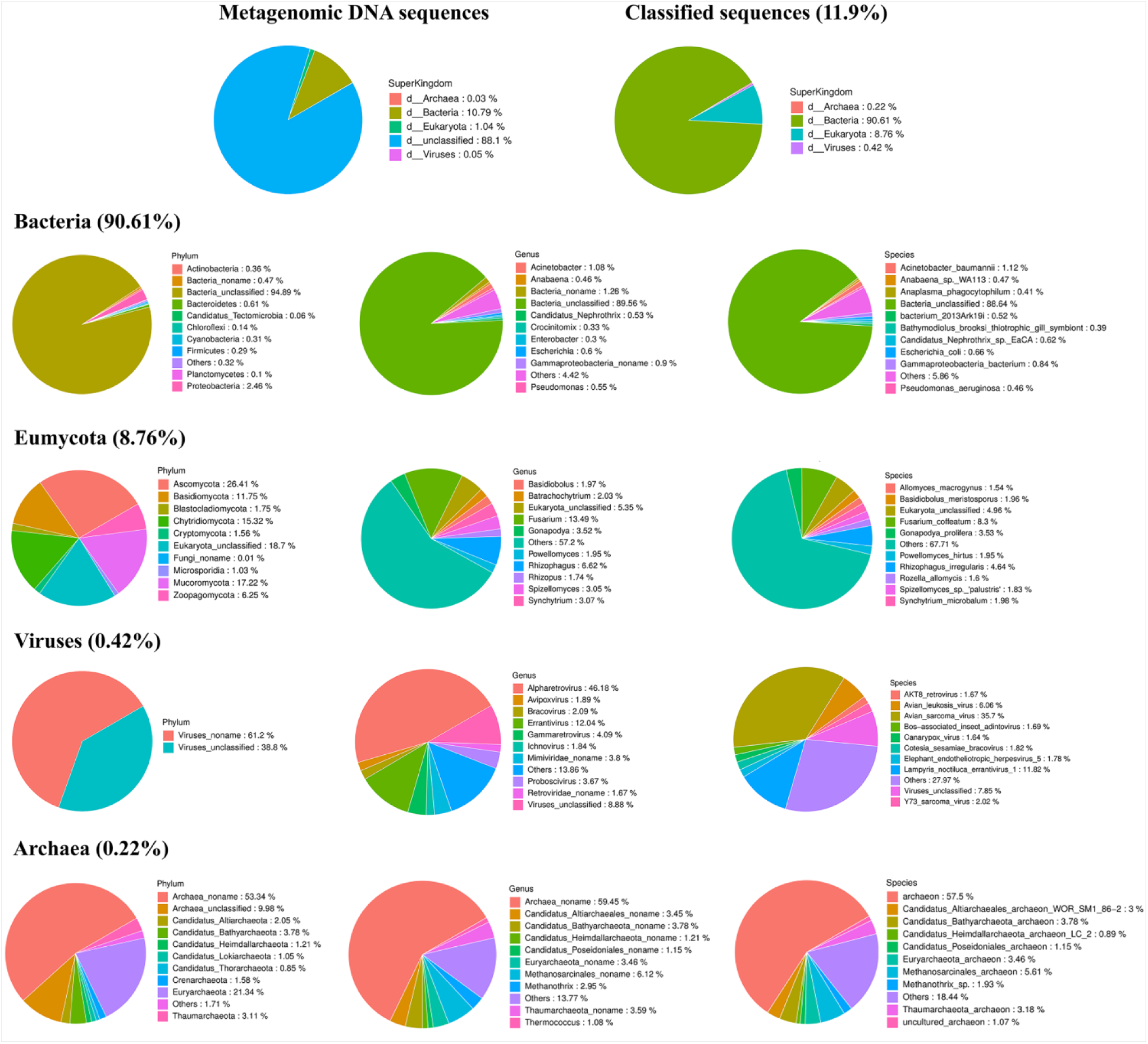
The microbial composition of the complex. Pie chart showing the composition of the microbial community of microbial complex. The proportions occupied by bacteria, fungi, viruses, and archaea in the microbial community are compared, as well as the proportions occupied by the phylum, genus, and species subordinate to each of the four kingdoms. In the bacterial kingdom, the pie charts from left to right represent the phylum, genus, and species taxonomic levels under the bacterial kingdom. The different colors in the pie chart for each taxonomic level represent different taxonomic categories. The size of the area occupied by the pie chart is the richness of the species at the phylum, genus, and species taxonomic level.

In summary, it suggested that the microbial complex contains a vast array of microorganisms, which include bacteria, fungi, virus, and archaea. They had provided a constant source of carbon for the whole community, assisting it in withstanding biotic and abiotic disturbances, and in the adaptation to the surrounding environment. Moreover, the microbial complex showed a strong and stable symbiosis.

### Functional analysis of the microbiome

The metagenomic results revealed a large quantity of unknown functional proteins in the microbial complex, which were mainly associated with amino acid transport and metabolism, carbohydrate transport and metabolism, gene replication and recombination, as well as repair, lipid transport and metabolism, and the synthetic transport and metabolism of secondary metabolites (Fig. S1A). This suggests that the microbial complex metabolism is highly active and provides enough raw material for the growth and development of bryozoans and other microorganisms. This is probably due to the abundant presence of Bacteroidetes in the microbial community, which can generate a large quantity of secondary metabolites and hydrolases, and that was also detected in the functional annotation of the genome. Many CAZymes were revealed in the microbial complex, among which the most dominant were glycosyl transferases (GTs), followed by glycoside hydrolases (GHs) and carbohydrate-binding modules (CBMs), with carbohydrate esterase (CEs) and polysaccharide Lyases (PLs) being the least abundant (Fig. S1). CAZymes hydrolyze many polysaccharides, such as microcrystalline cellulose, starch, xylem, chitosan etc., facilitating the use of nutrients in the environment by bryozoans and other microorganisms.

### Microbiome pathways within the microbial complex facilitate the use of nutrients by microorganisms

The top three functional categories in the KEGG annotation were ‘Global and overview maps’ (a top-level category), followed by carbohydrate metabolism and amino acid metabolism. (Fig. 4A). The enzymes involved in these pathways were also annotated in the genome (Fig. 4B), including phosphoenolpyruvate kinase (E4.1.1.32), which exploits the glycolytic and gluconeogenic pathways of carbohydrates, and citrate synthase (CS) (E2.3.3.1), isocitrate dehydrogenase (E1.1.1.42), and α-ketoglutarate dehydrogenase (E1.2.4.2), which are three key enzymes of the energy-producing tricarboxylic acid cycle. Through their activity, glucose can be converted into a variety of intermediate products and energy to be used for the growth and development of organisms and for the performance of all activities [23]. However, carbohydrates are rarely found in nature as monosaccharides, so CAZymes are necessary to perform hydrolytic role. Complex carbohydrates can be degraded to monosaccharides by CAZymes [24]. Glucose can be converted to pyruvate by the glycolytic pathway, and pyruvate can be transformed into acetyl coenzyme A by pyruvate dehydrogenase (E1.2.4.1). Acetyl coenzyme A combines with oxaloacetate in the presence of CS (E2.3.3.1) to form citric acid (Fig. 4B); it then enters the tricarboxylic acid cycle to generate energy for the growth and development of the organism. The amino acid metabolism follows that of carbohydrates (Fig. 4A), because some intermediates of other metabolic pathways are generated during the tricarboxylic acid cycle, such as acetyl coenzyme A, ferredoxin acid, and α-ketoglutarate (Fig. 4B).

**Fig. 4.**
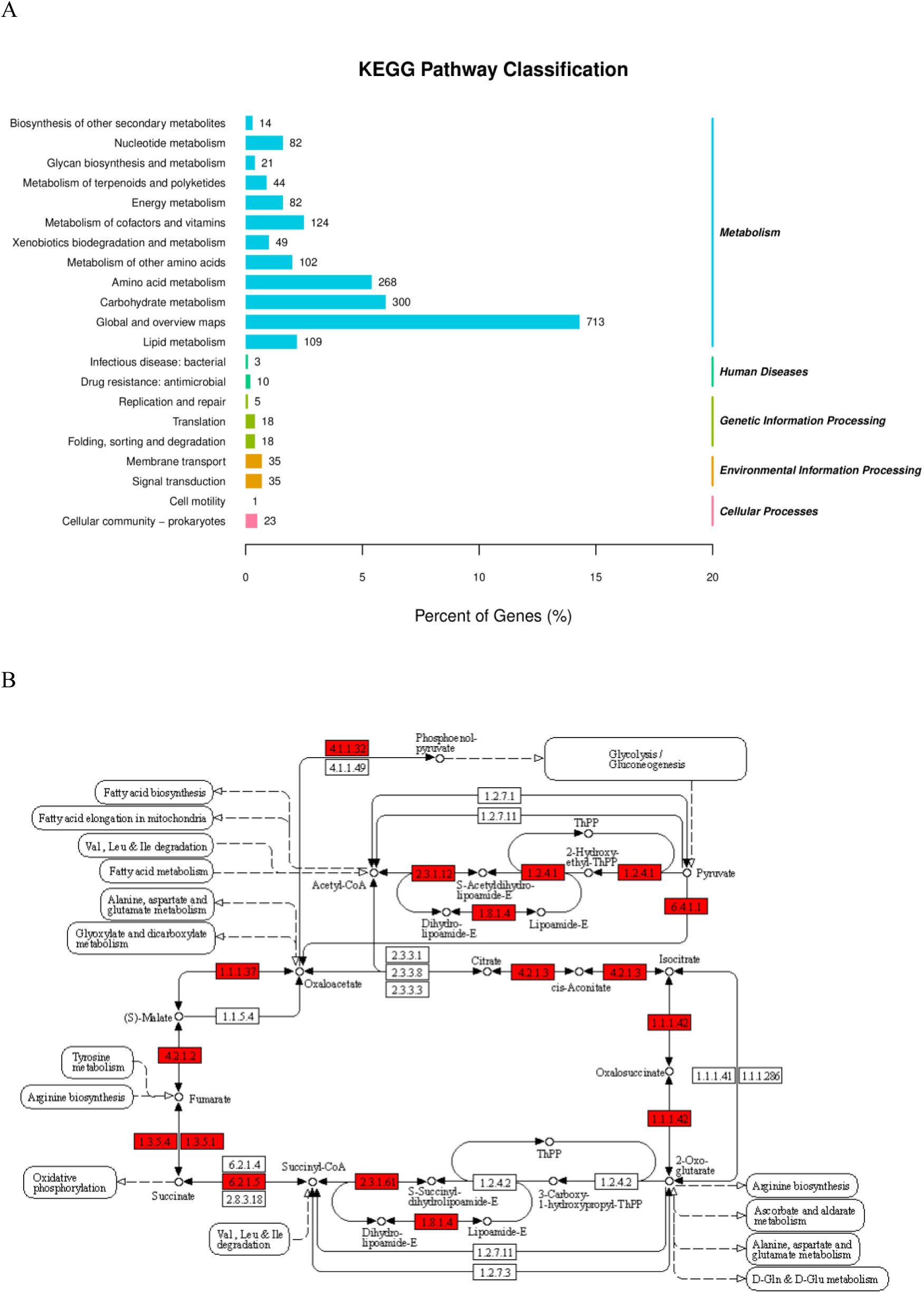
Annotation of the KEGG pathway in microbial macrogenomes. (A) KEGG pathway. The different colours are the different KEGG pathways. The size of the separate bar graph areas represents the abundance of the relevant pathways. (B) Citrate Cycle involved in KEGG enzymes from macrogenomic gene annotation. Rectangular boxes indicate the enzymes that catalyse the reaction (red border: the KO-related enzyme corresponding to the gene on the current data match), the box number is the EC number; Circles indicate metabolites (reactants or products of enzymatic reactions); The direction of the solid line arrow indicates the direction of the enzymatic reaction; The dashed arrows indicate that this product can then be related to other metabolic pathways via intermediate products; Rounded rectangular boxes represent other metabolic pathways.

### Genomic data indicate that microbial communities can resist abiotic stress

Genes with antibiotic effects (*tetA, bcrA, patB, msbA, lmrD*, *tet(X3)*, *poxtA*, *ermV*, *efrB, novA*, and others) were detected in the genome of the microbial complex (Fig. S2). Resistance genes, such as *tetA, bcrA, patB*, can inactivate antibiotics by increasing the efflux of antibiotic drugs, such as tetracycline, fluoroquinolone, cephalosporin, which may reduce the antibiotic-induced damage to the organism. Because of the presence of antibiotic-resistant genes, this microbial community is likely to have a considerably stronger ability to adapt to the environment, and to resist to antibiotics present in the water and other adverse conditions.

### The reduced virulence of pathogenic microorganisms contributes to maintaining the overall stability of the microbial complex

In the present study, various pathogenic fungi, such as *Magnaporthe oryzae*, *Aspergillus fumigatus*, and *Fusarium graminearum*, were detected within the bryozoan-dominated microbial complex. Notably, it implied that, when these pathogens became members of the complex, approximately one half showed a decline in virulence, and one fifth to one sixth completely lost their virulence (Fig. S3), which based on metagenomic detection of virulence factor genes (VFDB database) and literature-based annotation of known pathogenicity genes (PHI database). It was likely to suggest that the risk posed by the pathogens to other symbiotic microorganisms is minimal, and that it contributes to the maintenance of a healthy microbial complex.

### The metabolic production of microorganisms indicates their contribution to the overall microbial complex

Results of metabolomic assays showed that a total of eighty-one metabolites were annotated, and the ten highest-ranking were L-glutamate, L-glutamine, pyrrole-2-carboxylic acid, guanosine, betaine, L-methionine S-oxide, myo-inositol, spermidine, uracil, and phosphory lcholine, in this order (Fig. S4). The presence of glutamate-as well as glutamine-in large amounts, facilitates the growth and development of the host, and the response to abiotic stresses presents in the environment. Pyrrole-2-carboxylic acid, which is derived from the oxidation of hydroxy-L-proline, catalyzed by L-amino acid oxidase, display anti-inflammatory, analgesic, and anti-pyretic activities [25], that may assist biological organisms in the reaction against the invasion of pathogenic bacteria.

### Correlation between microbes and metabolites in the microbial complex

The present study showed that different metabolites were significantly relevant to most microorganisms within the complex (|r|> 0.50). A network map of microbe-metabolite interactions was also constructed by combining 16S functional predictions and metabolomics analysis. Interestingly, Firmicutes, Bacteroidetes, Planctomycetes, Cyanobacteria and Chloroflexi interact with the Cryptomycota, Zoopagomycota, Chytridiomycota, Blastocladiomycota and Mucoromycota phyla, which comprise fungi and viruses, to promote the production of L-lactic acid (M89T198), ketoleucine (M131T742) and adenine (M136T103), and the fungi (Cryptomycota, Mucoromycota, and Microsporidia) interact with bacteria (Chloroflexi, Candidatus Tectomicrobia), archaea and viruses to inhibit the production of cis-4-hydroxy-D-proline (M132T680) (Fig. 5). This is shown on the right side of Fig. 5, where the Acidobacteria, Firmicutes, Bacteroidetes and Proteobacteria phyla collaborate with the Zoopagomycota and Chytridiomycota fungal phyla to inhibit the production of 3-hydroxybenzyl alcohol glucoside (M124T793) (Fig. 5). The levels of this compound and cis-4-hydroxy-D-proline were greatly reduced due to the combined inhibition caused by a variety of microorganisms, thus, their presence was not identified in the top ten metabolites (Fig. S4). Ketoleucine and adenine are metabolites produced by a combination of microbial interactions, mainly through the conversion of glycine and glutamine, that was also annotated in significant amounts in the metabolomic analysis, and which indirectly supports the metabolomic findings. In addition, it was found that Ascomycota interact with Bacteroidetes to inhibit the production of prephenate (M227T567), urocanic acid (M139T104) and adenosine 2, 3-cyclic phosphate (M352T162) (Fig.5). These phenomena further suggest that the microorganisms in the whole community do not perform their function independently, but work in collaboration with each other closely. The results also implied that they coadjust to each other by secreting metabolites that regulate mutual inhibition or mutual promotion processes.

**Fig. 5.**
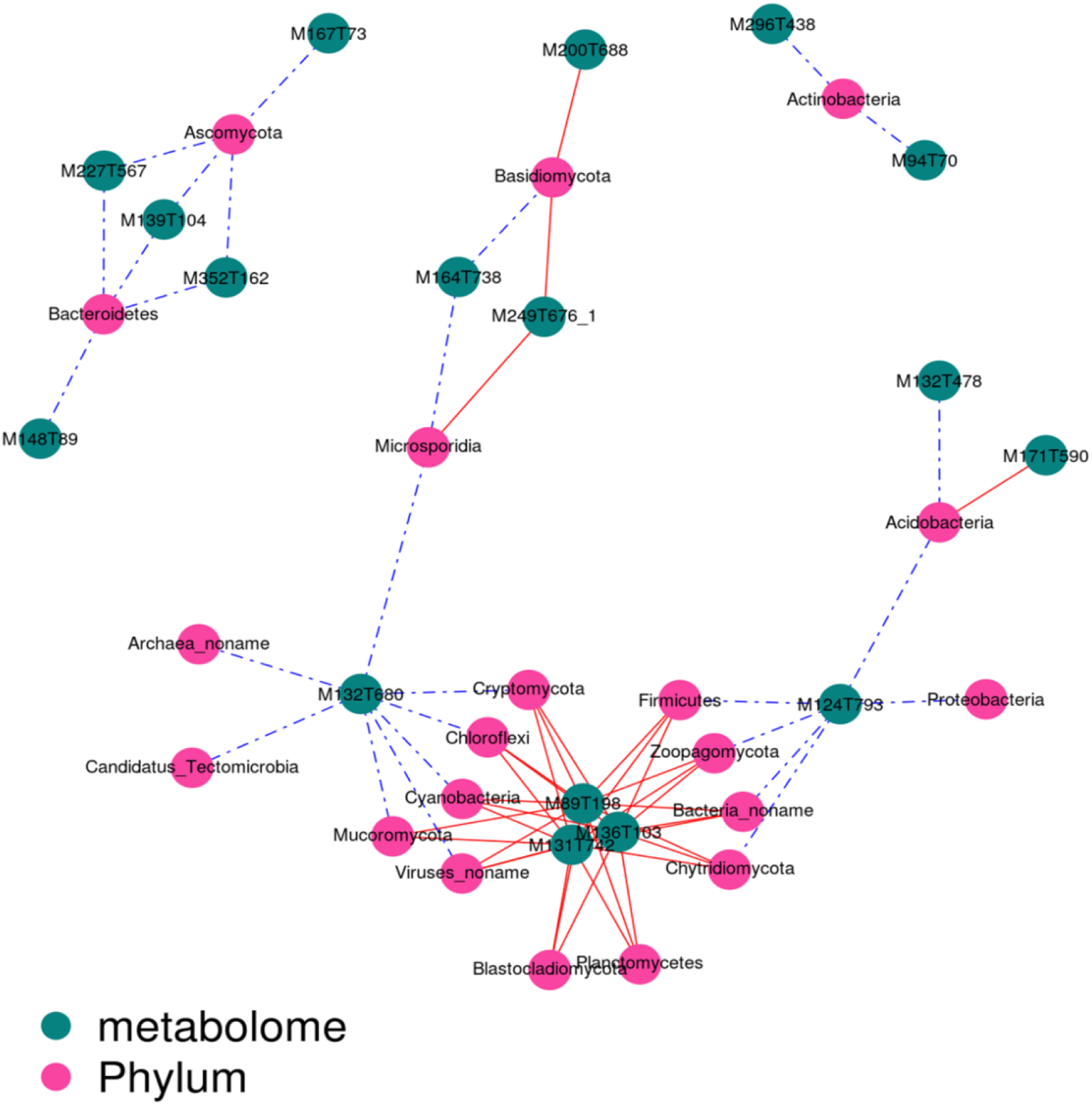
Correlation network diagrams represent the correlation values between phylum and metabolites. The red dots indicate phylum. Green dots indicate metabolites. The solid lines between phylum and metabolites indicate positive correlations and the dashed lines indicate negative correlations. Codes beginning with the letter M are for metabolites, they correspond to the metabolites in Table S3.

## Discussion

Bryozoans, a phylum of colonial, filter-feeding aquatic invertebrates [1,5,26]), which are increasingly recognized as apromising model for studying the combined, and more diverse and widespread-occurring from poles to tropics and from shallow coasts to the deep sea [27]. Many species are habitat-forming organisms that contribute to sediment production, shoreline protection and even biotechnological applications through the production of bioactive products [18,28]. The bryozoan species *P. magnifica* can affect water-taking buildings, causing serious damage to irrigation and other water supply systems [29](Geiser 1937). Moreover, they can affect the growth and development of indigenous organisms [9,30](. In addition, some bryozoan species and their symbiotic microorganisms produce the substance known as bryostatin, which has important medicinal properties that can inhibit the malignant growth of human tumors. Studies have reported that the bryostatin produced by symbiotic Proteobacteria, which has been widely used to combat cancer, has shown impressive therapeutic results in recent years [17]. Therefore, understanding the composition and function of bryozoan-dominated microbial communities is of great significance.

Microorganisms are ubiquitous [31]. They not only act as decomposers, but also increase productivity in the whole ecosystem through the autotrophic effect [32]. They are often not present individually, but rather as groups of various microorganisms that function together under specific environmental conditions [33]. These microorganisms are interdependent and mutually symbiotic. They play a crucial role in the decomposition and absorption of food, toxic substances, and even in medical treatments or the utilization of inorganic substances [34], thus benefiting the growth and development of symbiotic organisms. As microbiome research continues to evolve, high-throughput sequencing methods have allowed a more comprehensive understanding of microbial communities [33,35]. Understanding the nature of the symbiont-host relationship in terms of stability, mechanism, and benefits to both partners is a primary goal of symbiosis research [36]. The present study explored the composition of a unique microbial complex from a microscopic perspective. The results showed that the microbial communities established around bryozoans consisted of bacteria, fungi, viruses, and archaea, which could coexist to achieve a stable relationship, and form a microbial complex with a regular shape and texture.

Species annotations showed a vast array of Proteobacteria, Bacteroidetes, and Actinobacteria in the microbial community. Proteobacteria is one of the most diverse microbial phyla on Earth, and it is predominant in most biota. These bacteria contain nitrogen-fixing genes for gaseous nitrogen fixation [37], and ATP citrate lyase (*aclBA*), a key indicator or gene for the rTCA cycle, which was originally thought to be a potential CO_2_ fixation pathway [38]. Ribulose-1,5-bisphosphate carboxylase oxygenase and sulfur oxidation genes are found in Gammaproteobacteria [39]. This suggests that Proteobacteria can metabolize various forms of reduced and oxidized carbon, nitrogen and sulphur compounds; a process that has important implications for the formation and cycling of carbon, nitrogen, and sulphur in the habitat. Furthermore, previous studies have shown that Proteobacteria in symbiosis with bryozoans can produce bryostatin, which has a deterrent effect on omnivorous starfish and a repellent effect on the amphipod *Cheirimedon femoratus* [40,41]. This indicates that this bacterial phylum might have contributed to carbon and nitrogen fixation in bryozoans, and to be a source of defensive or aggressive substances that can protect the host from predators.

In addition to Proteobacteria, large quantities of Bacteroidetes were detected in the present study. This is considered as the most abundant bacterial group besides Proteobacteria and Cyanobacteria [42], which is in line with the results of our experiments. Bacteroidetes can secrete different carbohydrate-degrading enzymes [42]. In this study, the presence of a significant number of carbohydrate-active enzymes was also analyzed. The most abundant genes expressed in the genome were GTs, followed by GHs and CBMs. GTs catalyze the formation of glycosidic bonds in organisms [43–45], which contribute to the synthesis of glycoproteins on the cell membrane surface. This process is essential for the life activities of all organisms in the bryozoan-dominated microbial complex. GHs have the opposite function to GTs, as they break the glycosidic bonds of complex carbohydrates-such as cellulose and tributyrin-and hydrolyze them into monosaccharides [46,47]. The function of CBMs differs from that of GHs and GTs in that they have no catalytic activity. However, they can facilitate the binding of other enzymes to water-insoluble substrates, and assist carbohydrate-active enzymes in performing their functions [48]. Large amounts of carbohydrate-active enzymes can effectively degrade many different polysaccharides, including starch, xylan, cellulose, pectin, fucoidan, and mannan [49]. Some experimental results have shown that rhamnogalacturonan II is the most complex pectic polysaccharide. Even though this polymer contains 21 different glycosidic bonds, it can be degraded by Bacteroidetes [50]. It has also been reported that Bacteroidetes can regulate the antagonism among different bacteria to reduce bacterial competition for nutrients through a subtype of the type VI secretion system (T6SS)[51,52]. This suggests that Bacteroidetes plays an essential role in the degradation of carbohydrates, and even in the carbon cycle of the habitat, as well as in the harmonious symbiosis of microbial communities.

Based on the metagenomic results, thousands of eukaryota and archaea were also annotated in the bryozoan-dominated microbial complex, with eukaryota mainly detected in the Mucoromycota, Chytridiomycota, and Ascomycota phyla. Mucoromycota can generate carotenoids, as well as long-chain polyunsaturated fatty acids which are an integral part of cell membranes, and the immune system can quickly and effectively eliminate invasions of disease-causing organisms by relying heavily on intercellular communication [53]. The presence of polyunsaturated fatty acids facilitates the integrity of cell membranes, which probably helps to enhance the immune response of living organisms [54,55]. Moreover, it has also been reported that polyunsaturated fatty acids have an advantageous antibacterial effect [53]. In addition to the damage caused by pathogenic bacteria to the entire organism, abiotic factors-such as ultraviolet light, ozone, and exogenous free radicals-can also damage the bryozoan DNA and cell membranes, and cause protein denaturation, and lipid peroxidation [55]. Mucoromycota can produce large amounts of carotenoids, which have good antioxidant properties and are used to scavenge free radicals in advance, which may contribute to the protection of the organism. In addition to Mucoromycota, the Chytridiomycota and Ascomycota phyla also play a significant role within the bryozoan-dominated microbial complex. Chytridiomycota can utilize cellulose, a compound that most organisms find difficult to use [56]. This may contribute to reducing the competition for nutrients. Ascomycota can generate secondary metabolites with antioxidant and anti-inflammatory properties, such as dichloroisocoumarins [57], assisting the microbial complex to withstand both biotic and abiotic stresses, and to maintain the harmony and stability of the whole complex, as in the case of Mucoromycota.

Archaea live mostly in extreme environments; they can fix carbon and reduce mercury, which is particularly important for bioremediation [58]. In recent years, the industrial and agricultural development has caused widespread pollution of soil, environment, and it has degraded water quality. Many industrial effluents and pesticides are characterized by high salinity, high heat, strong acids, strong bases, and heavy metals [58]. Archaea can survive in extreme environments, degrade hydrocarbons, and reduce mercury present in the environment to volatile zero-valent mercury [59]. Our results suggest that the presence of eukaryota and archaea is likely to help bryozoans to cope with increasingly harsh environmental conditions. They have played a critical role in removing pollutants from water environments, consequently avoiding potential damage to bryozoan organisms.

Viruses are infective agents that can only parasitize other organisms to complete their reproduction cycle. The interaction between viruses and other organisms has been highly controversial, and the root of the controversy lies in the fact that all viruses are parasitic [60]. Since the discovery of the tobacco mosaic virus in the second half of the 19th century, there has been an effort to elucidate the pathogenic mechanisms of viruses in order to overcome viral diseases [61]. However, it has recently been reported that the coexistence between viruses and organisms might be beneficial. Viruses are widespread in the biota (a conservative estimate reports about 1031 species) [62], and can infect all animals ranging from polypores to cnidarians, and from bilaterians to chordates [60]. From the origins of animal evolution, viruses have interacted with microbial communities, and this symbiotic relationship allows them to share resources such as metabolites or genes. In the human body, 8% of the genes are composed of endogenous proviral forms of retroviruses [61]; and because the expression of endogenous retroviral syncytium promotes placental development, it reduces maternal rejection of the fetus, to a certain extent [60]. Pathogenesis also does not seem to be as common as symbiosis, or reciprocity, between viruses and organisms [63,64]. Notably, in the present study, many pathogenic microorganisms show a significant reduction in virulence or even loss of virulence. The potential mechanism behind this phenomenon may be related to the interaction of secondary metabolites secreted by other microorganisms or by bryozoans interacted with pathogenic microorganisms, and it should be further investigated.

The present study shows that bacteria, fungi, archaea and viruses live in the microbial complex, and the results of metabolomics analysis implied that they do not exist independently. A correlation diagram of microbe-metabolite interactions revealed that a variety of microorganisms work together to promote or inhibit the production of certain metabolites, such as L-lactic acid, ketoleucine, adenine, and cis-4-hydroxy-D-proline. This reflects the highly collaborative nature of the whole group: as the microorganisms interact with each other, they contribute to the production of specific metabolites and fulfill their role in the group.

The interaction between the dominant organism and microorganisms is generated through the secretion of metabolites, which have a profound effect on the growth and development of the dominant organism and on immune regulation [65]. Our results show the presence of large amounts of L-glutamate, L-glutamine, and pyrrole-2-carboxylic acid in the metabolites. This may be due to the abundant amino acid metabolic pathways in the microbial genome. L-glutamate, as a precursor of essential amino acids, can be converted to proline, glutathione antioxidant, and to glutamine via glutamine synthetase, to produce intermediate products of the tricarboxylic acid cycle and participate in the synthesis of nucleotides [66]. Besides, glutamate can also act as an external signal to increase the activity of antioxidant enzymes in the host, thereby counteracting the adverse effects of reactive oxygen species generated in the bryozoan under low-temperature conditions [67,68]. Therefore, the presence of glutamate as well as glutamine in large amounts may facilitate the growth and development of the microbial complex, and also the response to abiotic stresses in the environment. Pyrrole-2-carboxylic acid has a good bactericidal effect and reduces the invasions of pathogenic bacteria on microbial complexes.

In conclusion, the present study investigated the microbial composition, functions, and metabolites of a bryozoan-dominated microbial complex, and revealed a collaborative interaction with symbiotic microorganisms. All the members in the complex play their respective roles and work together to maintain the functionality of the microbial community. Because of their strongly collaborative group relationship, the community can successfully organize a new microbial ecosystem, display the shape and structure of a regular microbial complex, and even continuously increase in size. Also, a decrease or even a loss in virulence that occurs when pathogenic microorganisms become members of the complex, have been here reported. Besides, the results revealed the presence of a vast array of Proteobacteria, which have been previously reported in the literature to secrete bryostatins in response to bryozoan stimulation. This finding provides an important basis for future studies on bryostatin.

## Conclusions

The present study involves interdisciplinary research, which focus on one of the most ancient creature bryozoans, and the diversity analysis, metagenomic function analysis and metabolome analysis of aquatic microorganisms were covered. The results revealed the presence of a strong collaborative interaction with symbiotic microorganisms in the unusual bryozoan-dominated microbial complex. All the members in the complex play their respective roles and work together to maintain the functionality of the microbial community. Because of their collaborative group relationship, the community can successfully organize a new microbial ecosystem, display the shape and structure of a regular microbial complex, and even continuously increase in size. In addition, the results revealed the presence of a vast array of Proteobacteria.

## Limitations and suggest future studies

The limitations of the present study: lack of functional validation for virulence reduction, single sampling location and the number of samples, and no direct detection of bryostatin. Thus, it will be suggested that future studies include: (i) Identify the virulence genes of the advantageous species and analyze their expression levels to verify the changes in their virulence. (ii) Expand the sampling locations and increase the number of samples, and explore the common characteristics of the bryozoan-associated microbial complex. (iii) Analyze the artificial cultivation conditions for the bryozoans *P. magnifica*, extract the bryostatin and further conduct in-depth analysis of production mechanism for bryostatin.

## Declarations

### – Ethics approval and consent to participate

This research is solely focused on environmental microorganisms, and therefore does not involve any human or animal subjects.

### – Consent for publication

Not applicable

### – Availability of data and materials

Data and materials are available from the Ethics Committee of Hanshan Normal University (contact via Hanshan Normal University) for researchers who meet the criteria for access to confidential data.

## – Acknowledgments

The authors would like to thank grants from Special Project for Promoting the Distribution of Scientific and Technological Achievements from Guangdong Province to Counties and Towns (2025B0202010049), Project for the Rural Science and Technology Commissioner Project of the Department of Science and Technology of Guangdong Province (KTP20240860).Guangdong Province university key field special grant (2022ZDZX4029, 2024ZDZX2036), Guangdong Provincial Key construction discipline research ability enhancement project (2021ZDJS041), Special Project of the “Double Hundred” Initiative (XSB202406), Guangdong Provincial Key Laboratory of Functional Substances in Medicinal Edible Resources and Healthcare Products (2021B1212040015). The authors acknowledge Guangzhou Jirui Gene Technology Co., LTD. for providing facility and essential materials during the experiment.

